# A new mouse model for a high-risk AMD variant, *ARMS2^A69S^*

**DOI:** 10.64898/2026.10.02.756275

**Authors:** Navdeep Gogna, Gayle B. Collin, Lillian F. Hyde, Lisa Stone, Ravi Pandey, Victoria E. Kennedy, Mark P. Krebs, Juergen K. Naggert, Patsy M. Nishina

## Abstract

Age-related macular degeneration (AMD) is the leading cause of irreversible central vision loss in older adults. The highest genetic risk for AMD maps to Chromosome 10q26. However, due to strong linkage disequilibrium between two genes, *ARMS2* and *HTRA1* located in this region, it has been difficult to determine whether one or both genes contribute to the AMD-disease risk. Functional studies of *ARMS2* have been hampered due to the lack of animal models, as orthologs are only found in the higher primates and the gene is absent in rodents. To model *ARMS2* biology *in vivo*, we used CRISPR-Cas9 to insert the complete human *ARMS2* gene carrying the A69S risk variant (rs10490924), into a region of homology between humans and mice, on mouse Chromosome 7, 3.5 kb upstream of *Htra1*, preserving its native promoter, spacing, and orientation. We also generated a mouse model bearing its wild-type counterpart *ARMS2*^*A69*^ (*ARMS2*^*WT*^). RNA analysis revealed tissue- and cell-type-specific patterns of *ARMS2* expression with enrichment in neural tissues, testis, and eye, and within the eye, in retinal horizontal cells and the retinal pigment epithelium (RPE). *ARMS2* expression was consistent with human expression data and unaltered by the A69S mutation. By 12 months of age, *ARMS2*^*A69S*^ mice developed AMD-relevant phenotypes including increased fundus spots, A69S-specific thinning of the outer nuclear layer, and reduced scotopic ERG amplitudes. Transcriptomic profiling of postnatal day 14 retina and RPE-enriched fractions revealed reduced mitochondrial oxidative-phosphorylation genes, corroborated by reduced retinal mtDNA copy number and lower maximal respiration, alongside preliminary results that suggest sphingolipid remodeling in the RPE. This humanized model links the *ARMS2* A69S variant to mitochondrial dysfunction and provides an *in vivo* platform for defining AMD mechanisms and testing therapies.

## Introduction

Age-related macular degeneration (AMD) is the leading cause of irreversible vision loss among the elderly in developed nations^1^ and a major cause of blindness worldwide^2^. AMD affects the macula, a specialized central region of the retina that is densely packed with cone photoreceptors and mediates high-acuity, color and daytime vision; as a result, central vision is affected, while the peripheral vision is retained^3^. Understanding AMD is challenging because it is a complex disease, in which onset and severity is shaped by many interacting genetic and environmental factors^1,4^. AMD is also heterogeneous, involving multiple disease mechanisms (complement dysregulation^5^, inflammation^6^, oxidative stress^7^, altered lipid metabolism^8^, retinal pigment epithelium (RPE) dysfunction^9^ and choroidal vascular changes^10^), and a broad phenotypic spectrum (dry [atrophic] versus wet [neovascular] AMD, and drusen subtypes including hard, soft, cuticular, and reticular pseudodrusen or subretinal drusenoid deposits^11-14^). Further, the primary site of disease initiation within the photoreceptor-RPE-Bruch’s membrane-choriocapillaris (PRBC) complex is a central, unresolved question. The drusen and basal deposits that accumulate at the RPE-Bruch’s membrane interface implicate the RPE^15^, yet choriocapillaris loss can be detected early and may precede overt RPE changes^16^. Because the PRBC layers are structurally and metabolically coupled, distinguishing a primary insult from secondary, downstream consequences has proven difficult. Compounding the problem, several of the strongest risk pathways: complement activation, oxidative stress and lipid metabolism, operate systemically, raising the possibility that AMD is driven, at least in part, by circulating or systemic factors rather than defects intrinsic to a single ocular cell type^17^. Together, these observations have framed AMD as a collection of related pathological processes that converge on the PRBC complex and not as a single homogeneous disease.

Genetic factors account for 46% to 71% of the variation and severity observed in AMD^18^. Genome-wide association studies have identified 52 independent common and rare variants across 34 loci, implicating pathways that include complement activation, lipid metabolism, and extracellular-matrix organization^19^. Among these, the Chromosome 10q26 locus has been most strongly associated with increased AMD risk^20,21^. However, the biological basis for this association has remained unresolved, largely because two genes - *Age-related maculopathy susceptibility 2* (*ARMS2*) and *High*-*temperature requirement protein A1* (*HTRA1*) lie in strong linkage disequilibrium in this region that couples their risk-associated variants, confounding statistical efforts to assign causality to either or both genes^22^. While *HTRA1* encodes a known serine protease^23^, the structure, function and localization of the predicted 11-kDa ARMS2 translation product is not firmly established. Further, the *ARMS2* gene is present only in higher primates^24^ with no ortholog in mice or other common laboratory animals. This absence has precluded conventional loss-of-function approaches and has left the field reliant on *in vitro* systems and patient-derived cells, which cannot fully recapitulate the aged retinal microenvironment in which AMD manifests. Prior attempts to study *ARMS2* using immortalized cells or ocular tissue have reported ARMS2 localizes to variable subcellular and extracellular components: in the mitochondria^20^, cytoplasm^25^ or in the extracellular matrix of the choroid pillars of the human eye^26^. *ARMS2* has also been reported to express in human blood-derived monocytes and in human-induced pluripotent stem cell-derived microglia (iPSdM)^27^.

Multiple *ARMS2* variants have been implicated in AMD risk, with the missense variant, rs10490924 (G>T:A69S)^20,25,26,28^, encoding an alanine-to-serine substitution at position 69 (A69S) as the most frequently observed *ARMS2* risk allele in the human population. The effect of this mutation on ARMS2 protein structure and function is unknown, although, one study reported that the A69S variation in ARMS2 creates a new putative phosphorylation site that disrupts a predicted α-helix^20^. Previously, two transgenic mouse models expressing human *ARMS2*^*A69S*^ and its control *ARMS2*^*A69*^ have been reported^29,30^. Neither model showed pathological changes in the eye. The constructs to generate these mouse models used a ubiquitous cytomegalovirus (CMV)^29^ or chicken actin (CAG) promoter^30^, and were either inserted into the ROSA locus^29^ or randomly in the mouse genome^30^. These findings suggested that the non-physiological overexpression of *ARMS2*^*A69S*^ was insufficient to recapitulate AMD-like features and underscored the need for the human *ARMS2* gene to be expressed at appropriate levels and in an appropriate spatial and temporal pattern.

In this study, we generated and characterized a mouse model in which the human *ARMS2* gene carrying the A69S risk variant (and its non-risk A69 counterpart) was introduced at a site upstream of the *Htra1* locus, in a region that is homologous between humans and mice, thus preserving genomic localization and orientation. Further, a native *ARMS2* promoter included in the construct preserved endogenous regulatory elements. Human *ARMS2* expression and its tissue and cellular localization within the eye were confirmed by qRT-PCR and RNAscope, respectively, and phenotypic changes were assessed by fundus imaging, optical coherence tomography (OCT), electroretinography (ERG), and -omics approaches. This model provides an *in vivo* platform to explore ARMS2-driven mechanisms of AMD-risk susceptibility and a system in which to test therapeutic strategies targeting these pathways.

## Materials and Methods

### Mice

Mouse strains used in this study - C57BL/6J-*Htra1*^*em1(ARMS2A69S)Pjn*^, C57BL/6J-Htra1^em1(ARMS2A69)Pjn^ and C57BL/6J (hereby referred to as *ARMS2*^*A69S*^, *ARMS2*^*WT*^ and B6, respectively), were bred and maintained under standardized conditions of 12:12 hour (h) light:dark cycle in the Research Animal Facility at The Jackson Laboratory (JAX). Mice were maintained in pressurized individual ventilation caging, provided with food (LabDiet 5K52) and HCl-acidified water (pH 2.8–3.2) *ad libitum* and monitored for the maintenance of a pathogen-free environment. Animal protocols were approved by the JAX Institutional Animal Care and Use Committee (AUS99089) and complied with guidelines set forth by the ARVO Animal Statement for the Use of Animals in Ophthalmic and Vision Research.

### Generation of humanized ARMS2 knock-in (KI)

CRISPR-Cas9 gene editing mediated by homology recombination was used to create a murine model bearing the human *ARMS2*^*A69S*^ AMD susceptibility allele (JAX Genomic Engineering Technology, The Jackson Laboratory, Bar Harbor, ME). Briefly, a donor plasmid (pUC57) containing the full length human *ARMS2* gene (including >2 kb upstream and the A69S variant sequence (rs10490924; c.205 G>T)) flanked by 5’ and 3’ homology arms, was co-injected with Cas9 protein with a target-specific sgRNA into fertilized B6 zygotes. The humanized *ARMS2* transgene inserted into the homologous region of mouse Chromosome 7, approximately 3.5 kb upstream of *Htra1*. KI founders were identified by long range PCR, backcrossed to B6 and *ARMS2* KI carriers were intercrossed.

Subsequently, the *ARMS2*^*WT*^ mouse model was generated by reverting the A69S mutation in *ARMS2*^*A69S*^ mice back to the wild-type (A69) allele by CRISPR-Cas9 HDR-mediated gene editing using a single stranded donor oligonucleotide. Briefly, two crRNAs targeting the region flanking the A69S variant (ARMS2_WT_crRNA1: 5′-GATTTTAGAAGCTGGGATCA-3′; ARMS2_WT_crRNA2: 5′-CAGTGTGGATTTTAGAAGCT-3′), were combined with Cas9 protein to form ribonucleoprotein complexes and co-injected with a single-stranded donor oligonucleotide encoding the wild-type A69 sequence into fertilized (*ARMS2*^*A69*^ X B6) zygotes. A69 founders were backcrossed to B6 and resulting F1 progeny were intercrossed to generate mice for downstream studies.

### Genotyping and sequencing

The mice were genotyped to confirm the presence of the humanized *ARMS2* gene in the mouse genome. Genomic DNA was extracted from tail tips (*<*2 mm) or ear punches using a standard protocol^31^. Briefly, the tissues were incubated in 50 mM sodium hydroxide solution for 30-50 min at 95°C, followed by the addition of 1M Tris HCl neutralizing reagent buffer (final concentration 200 mM), vortexed and centrifuged at 3600 rpm for 8 min at 4°C. For PCR amplification,1 μl of the supernatant was used for genotyping. To determine the absence/presence of the *ARMS2* transgene, we used the following primer combinations: F-5’-TTAGCTGCTGACTGCCACTC-3’ and R -5’-TATGGCCCCGTGTGGATGTC-3’ (WT allele; absence of *ARMS2*) and F-5’-TGGCTGCTCCATTCCTATTC-3’ and R-5’-GAATGTGTACCCACCCAAGG-3’ (Tg allele; presence of *ARMS2*). To determine if the inserted sequence was *ARMS2*^*WT*^ or *ARMS2*^*A69S*^, PCR was performed using primers: F-5’-CTGGCTGAGTGAG-3’, R-5’-GCAGGTGCATTGG-3’ and products subsequently underwent Sanger sequencing (G, A69; T, A69S).

### qRT-PCR

The expression of *ARMS2* in different tissues was tested by qRT-PCR. RNA was isolated as described previously^32^ using a RNeasy Kit (QIAGEN, Germantown, MD, USA). cDNA was then synthesized using the Superscript IV First Strand cDNA Synthesis kit (ThermoFisher Scientific, Waltham, MA, USA) and qRT-PCR was performed with iTaq Universal SYBR Green SuperMix (Bio-Rad, Hercules, CA, USA), both using an established manufacturer’s protocol. For qRT-PCR, we used gene-specific primers (ARMS2: F - 5’-TCGGTGGTTCCTGTGTCCTTCATT-3’ and R - 5’-TCACCTTGCTGCAGTGTGGATGAT-3’ and mActb: F - 5’ - CCAGTTCGCCATGGATGACGATAT-3’ and R – 5’ - GTCAGGATACCTCTCTTGCTCTG-3’) and a CFX96 Real-Time PCR Detection system (Bio-Rad). The comparative CT method (ΔΔCT) was applied to calculate a relative-fold change in transcripts and quantified using 2^−^ΔΔCT with β-actin as an internal calibrator.

### Fundus imaging

Mice were anesthetized with isoflurane provided by a vaporizer (Kent Scientific, Torrington, CT, USA) and pupils dilated using 1 % cyclopentolate or 1 % atropine (Akorn, Laki Forest, IL, USA) eye drops. Fundus images were acquired using a Micron IV fundus camera (Phoenix Research Laboratories, Pleasanton, CA, USA) and processed using a protocol described previously^33^. Images were analyzed in Fiji using the Trainable WeKa Segmentation plugin, a supervised pixel-classification tool that combines image-derived features with a machine-learning classifier. A classifier was trained on a representative subset of images chosen to span the range of spot intensities, size, and variation in disease severity present in the full dataset. Pixels were assigned to three classes - “spot”, “background” or “blood vessels”. The area around the optic disc was excluded in the analysis. Classification used the default FastRandomForest classifier (Weka^34^) with 200 trees. Annotations were added iteratively and the classifier retrained until visual inspection confirmed accurate separation of spots from background. The trained classifier was then applied, unchanged, to every image in the dataset to generate per-pixel probability maps for the “spot” class. Spots were counted and measured with the Analyze Particles command, restricting detections to objects of size (pixel^2^) 50 – infinity and circularity (0.0–1.0) to exclude noise and non-spot structures. For each image, the number of spots and percentage spot area were recorded.

### Optical coherence tomography (OCT)

Ocular biometry was determined by OCT using a Bioptigen Ultrahigh-resolution (UHR) Envisu R2210 SD-OCT imaging system (Leica Microsystems, Buffalo Grove, IL, USA). Briefly, mice were anesthetized with an intraperitoneal (i.p.) injection of xylazine (80 mg/kg) and ketamine (16 mg/kg) in normal saline. A drop of GenTeal Severe gel (Alcon, Fort Worth, TX, USA) was applied to both eyes of mice to prevent corneal drying prior to image acquisition. Volume scans (1000 × 100 × 10, 1.9 mm diameter) were obtained, and images were processed in Fiji using methods described previously that utilized the long posterior ciliary arteries as landmarks to orient the eyes^33,35^. Briefly, to determine retinal layer thicknesses, datasets were transformed orthogonally and oriented rotationally based on choroidal vascular landmarks. Pixel intensities were obtained from a z-profile of circular ROIs positioned superior and inferior to the optic nerve head, 400 µm from its center. Approximate intensity peaks and valleys selected manually along this z-profile were repositioned precisely using an Excel template, which defined midpoints along each layer boundary where intensity was half the local peak-to-valley difference and calculated layer thickness as the distance between consecutive boundary midpoints.

### Electroretinography (ERG)

Eyes were dark adapted and then anesthetized with an i.p. injection of ketamine/xylazine mixture as described above. Additional anesthetic was given if akinesia was inadequate. The equipment and protocol used here is as previously described^36^. Briefly, dark-adapted, rod-mediated ERGs were recorded with responses to short-wavelength flashes over 4.0-log units to the maximum intensity by the photopic stimulator. The signals were sampled at 1-ms intervals and averaged.

### RNAscope

Eyes were enucleated, embedded and oriented (for dorsal/ventral cutting) in optimal cutting temperature compound (OCT; Tissue-Tek), and snap-frozen in isopentane on dry ice. Ten-micron thick cryosections were cut on a cryostat, set at 15°C, collected on SuperFrost Plus slides (ThermoFisher, Waltham, MA, USA) and stored at −80°C until use. *In situ* hybridization was performed using the RNAscope Multiplex Fluorescent Reagent Kit v2 (Advanced Cell Diagnostics (ACD, Newark, CA, USA) following the manufacturer’s protocol for fresh-frozen tissue. Sections were removed from −80°C, equilibrated at room temperature, and fixed in pre-chilled 4% paraformaldehyde in 1X PBS for 15 min at 4°C. Sections were then dehydrated through an ascending ethanol series (50%, 70%, 100%, 100%) and air-dried. A hydrophobic barrier was drawn around each section (ImmEdge pen), and sections were treated with RNAscope Hydrogen Peroxide for 10 min at room temperature, followed by RNAscope Protease IV for 15 min at room temperature, with 1X PBS washes between steps. Target probes (*Hs-ARMS2*-C1 and *Mm*-*Lhx1*-C2) were hybridized for 2 h at 40°C in a HybEZ II oven (ACD). Probe signals were amplified with reagents provided in the kit. Channel-specific signals were developed sequentially by incubating with the corresponding HRP (HRP-C1/C2), the assigned (Opal/TSA Vivid) fluorophore (520 for ARMS2 and 650 for Lhx1 at 1:800 dilution) for 30 min at 40°C, and HRP blocker, with wash-buffer rinses between steps. Nuclei were counterstained with DAPI, and sections were mounted with ProLong Gold Antifade/ Vectashield (Vector Laboratories, Newark, CA, USA). Positive control (3-plex; ACD) and negative control (dapB; ACD) probes were run in parallel on adjacent sections to confirm RNA integrity and assess background. Fluorescent signals were visualized using an Axio Observer.z1 fluorescence microscope with an ApoTome 2.0 accessory (Carl Zeiss, Oberkochen, Germany) and images processed using Fiji software^37^.

### Transcriptomic analysis

Enriched fractions of retina and RPE were obtained from 14-day old mice, using a protocol adapted from Wang et al. ^38^. Eyes were enucleated after euthanizing the animals and eye dissections performed in a 100 mm dish containing 1× PBS with 10 mM ribonucleoside vanadyl complex (New England BioLabs, Ipswich, MA), on ice. After removing the anterior eye and lens, the posterior eyecups were incubated in hyaluronidase for 10 min at 37°C. Retinas were separated from the RPE-choroid/sclera (RPE-CS) and placed in RNAlater (Qiagen, Hilden, DE). The remaining RPE-CS complex was incubated in RNAprotect Cell Reagent (Qiagen, Hilden, DE) on a shaking platform for 3 min at room temperature. To separate the RPE and CS, the tissue was gently withdrawn from the liquid with forceps, causing the RPE sheet to detach visibly at the liquid–air interface and release back into the RNAprotect solution. The RPE sheet (in RNAprotect) was snap-frozen on dry ice and immediately stored at −80°C. Retina and choroid/sclera samples were stored overnight at 4°C and then moved to −80°C the next day. Bulk RNA-seq (NovaSeq X Plus, 150-bp paired-end; KAPA mRNA HyperPrep for retina and SMARTer Stranded Total RNA-Seq for RPE) were performed as described^39,40^. Gene-level read counts were analyzed for retina and RPE samples from female *ARMS2*^*A69S*^ and B6 mice. Mitochondrial genome–encoded genes were identified using Ensembl annotations. Mitochondrial rRNA was excluded from the mitochondrial fraction because its abundance differed between tissues. Cross-tissue contamination was assessed using marker panels - RPE markers (*Rpe65, Best1, Ttr, Rgr, Lrat, Tyr, Mlana*) in retina and photoreceptor markers (*Rho, Sag, Pde6b, Pde6g, Gnat1, Nrl, Rp1, Prph2, Rbp3, Cnga1, Rom1, Pdc*) in RPE - with contamination fraction estimated as the ratio of each marker’s expression in the recipient tissue to its expression in the source tissue. The RPE contamination in the retina was subtracted from the observed counts before differential expression analysis, which was performed using PyDESeq2 v0.5.4 with genotype as the sole design factor, comparing *ARMS2* with B6J. Genes with at least 10 counts in at least three libraries were retained, yielding 26,536 genes in retina and 24,041 in RPE. Differential expression was assessed using Wald tests, with significance defined as a Benjamini–Hochberg adjusted *p* < 0.05. When multiple Ensemble identifiers mapped to the same gene symbol, the entry with the largest absolute Wald statistic was retained for downstream analysis. Preranked gene set enrichment analysis was performed using GSEApy, with genes ranked by the Wald statistic. Each gene set collection was analyzed separately using 1,000 permutations. Gene sets containing 10–500 genes were included for the Hallmark collection and 4– 500 genes for curated collections. Normalized enrichment scores were reported, and significant enrichment was defined as a false discovery rate (FDR) q-value < 0.05, calculated within each collection.

### Mitochondrial quantification

Relative mitochondrial DNA (mtDNA) copy number was determined by quantitative PCR (qPCR) as the ratio of a mitochondrial amplicon to a single-copy nuclear reference amplicon measured from the same lysate. The mitochondrial target was amplified within a region selected to avoid co-amplification of nuclear mitochondrial DNA segments (NUMTs). Primer sequences used: mtDNA non-NUMT, [F-5′-CTAGAAACCCCGAAACCAAA-3′; R: 5′-CCAGCTATCACCAAGCTCGT-3′]; n-*TERT*, [F-5′-CTAGCTCATGTGTCAAGACCCTCTT-3′; R-5′-GCCAGCACGTTTCTCTCGTT-3′]. Retinas were isolated from enucleated eyes in ice-cold DPBS and suspended in alkaline lysis buffer (25 mM NaOH, 0.2 mM EDTA, pH ~12), incubated for 30 min at 95°C to release and denature total DNA and cooled to 4 °C. Lysates were neutralized by addition of 40 mM Tris-HCl (pH 5.0) and either used immediately or stored at −20°C. Crude lysates were diluted to have a starting amount of 10 ng, and qPCR reaction run using established manufacturer’s protocol with SYBR Green SuperMix (Bio-Rad, CA, USA). Each sample was run in technical duplicates, alongside no-template controls and a water blank carried through the entire workflow. Relative mtDNA copy number was calculated using the comparative Ct method. Within each sample, ΔCt was defined as Ct(*TERT*) − Ct(mtDNA), and fold change between mutant and control was calculated as 2^(ΔΔCt).

### Seahorse Mito Stress Test assay

Retinal mitochondrial respiration was measured ex vivo using a Seahorse extracellular flux analyzer (Agilent, Santa Clara, CA, USA), adapted from the protocol described by Shetty et al^41^. Sensor cartridges were hydrated in XF calibrant overnight (12–14 h) in a non-CO_2_ incubator at 37°C. On the day of the assay, freshly enucleated eyes from euthanized mice were dissected to isolate the neural retina free of RPE; the retinal cup was flattened with four radial incisions, and 1 mm disks were cut with a biopsy punch from regions adjacent to the optic nerve head to maintain layer homogeneity. Each intact punch was placed separately in a well (with the ganglion cell layer facing up) of a Seahorse XFe96/XF Pro Cell Culture Microplate in XF DMEM (pH 7.4) supplemented with 12 mM glucose, 2 mM L-glutamine and 0.5 mM sodium pyruvate, and equilibrated for 45 min in a room-air (non-CO_2_) incubator at 37°C to allow CO_2_ equilibration between tissue and medium. Oxygen consumption rate (OCR) was measured using a modified XF Cell Mito Stress Test consisting of sequential injections of FCCP (0.5 µM) and rotenone/antimycin A (0.5 µM); the oligomycin injection was omitted owing to its toxicity to retinal punches^41^. Measurement cycles comprised 3 min mix, 2 min wait, and 5 min measure, with five baseline cycles, four cycles after FCCP, and four cycles after rotenone/antimycin A. Non-mitochondrial respiration was defined as the minimum OCR after rotenone/antimycin A; basal respiration, maximal respiration, and spare respiratory capacity were calculated in Wave software (v2.2, Agilent). At least 3-4 biological replicates per strain and 3 technical replicate punches per eye were analyzed, and punches yielding flat or non-responsive traces were excluded as non-viable. Each plate contained one biological replicate each from the two strains under study (*ARMS2*^*A69S*^ and *ARMS2*^*WT*^). The results were analyzed using JMP statistical analysis software (SAS Institute, Cary, NC, USA), using restricted maximum likelihood (REML) mixed models, with strain, sex, and their interaction as fixed effects and plate (nested within sex) as a random effect. Technical replicate wells were averaged to a single value per animal before analysis (n = 7 plates; 3 female, 4 male; one animal per strain per plate; 3 technical replicates per strain per plate).

### Mass Spectrometry Imaging (MSI)

Immediately after enucleating the eyes, an insoluble paint dot was placed on the dorsal globe for orientation of the eye. The eyes were then embedded in M-1 Embedding Matrix (Epredia, Kalamazoo, MI, USA), rapidly frozen on dry ice, and stored at −80°C until sectioning. For cryosectioning, the specimens were briefly maintained at −20°C prior to collect 10-µm sections on the conductive side of Intellislides (Bruker, Billerica, MA, USA) at -13 °C. A matrix (e.g. 1,5-diaminonaphthalene for positive ion mode) was applied to slides through sublimation using the HTX Sublimate as it improves the high spatial resolution of lipids (≤5 µm resolution). Slides were then analyzed using the Bruker timsTOF FLEX MALDI-MSI instrument in the JAX Mass Spectrometry and Protein Chemistry service laboratory as described previously^42^. All raw data was exported as .d files to the Bruker SCILS Pro MALDI-MSI software analysis package for data visualization and region of interest (ROI) interpretation of the data according to the MSPC workflow and commercial guidelines. Results were analyzed in SCiLS lab where lipid species were annotated from accurate mass and collision cross-section. Mean peak areas per RPE region of interest (ROI) were exported after per-pixel total-ion-count (TIC) normalization. n = 3/sex/strain were used for MSI. Annotations were screened for chemical plausibility in mammalian tissue before analysis. Non-mammalian classes (betaine lipids, DGTS; sulfoquinovosyl-diacylglycerols, SQDG) were removed. Acyl compositions were evaluated per chain: fatty acids were allowed up to six double bonds, four or more double bonds required a chain of at least 20 carbons, and sphingolipids were modelled as an 18-carbon sphingoid base with at most two double bonds plus one fatty acid. This retained very-long-chain polyunsaturated species characteristic of retinal tissue while excluding compositions such as SM 28:7 or HexCer 30:6 that cannot arise from mammalian fatty-acid biosynthesis. Curation removed 87 of 404 male and 111 of 446 female annotations. Remaining annotations sharing a lipid name and adduct were collapsed by summation, and features with intensity below 100 in more than half the sections within a sex were excluded, leaving 156 (male) and 184 (female) features. TIC-normalized intensities were log_2_-transformed. Pairwise group differences were tested by Welch’s two-sample t-test on log_2_ values with Benjamini-Hochberg false-discovery-rate (FDR) correction within each comparison and analysis level. Ordered trends across B6 - A69WT - A69S (scored 0, 1, 2) were tested: a linear contrast (ordinary least-squares slope, two-sided t-test) and the non-parametric Jonckheere–Terpstra (JT) test with exact two-sided p-values obtained by complete enumeration of label permutations (1,680 male; 4,200 female).

### Statistical Analysis

Statistical analysis for various analysis was performed in GraphPad Prism (version 11.1.0) and/or JMP (version 18.2.0) statistical analysis software. Effects of strains and sex and their interactions, as well as plates for Seahorse assay were noted.

## Results

### Generation of a new humanized *ARMS2*^*A69S*^ knock-in mouse model

To study the contribution of the *ARMS2* A69S risk variant to AMD-associated phenotypes in a mouse model, we introduced the full length human *ARMS2* gene (Chromosome 10q26) as a knock-in (KI) into the homologous region on mouse Chromosome 7 using CRISPR-Cas9 technology (Figure 1). A 6 kb human genomic fragment spanning the *ARMS2* 5’ regulatory region, and both *ARMS2* exons with the A69S substitution (rs10490924) in exon 1 was inserted 3.5 kb upstream of the mouse *Htra1* transcriptional start site, reproducing the spacing and orientation of *ARMS2* relative to *HTRA1* in the human genome. Homologous recombination was directed by 2.1 kb left and 2.4 kb right homology arms flanking the insert. Correctly targeted clones were identified by long-range PCR across each arm and confirmed by Sanger sequencing of the insert, which verified the A69S codon and the absence of off target mutations; founders transmitted the allele through the germline at expected Mendelian ratios (founder transmitting single-copy transgene to the next generation (F1) as 1:1).

**Figure 1:**
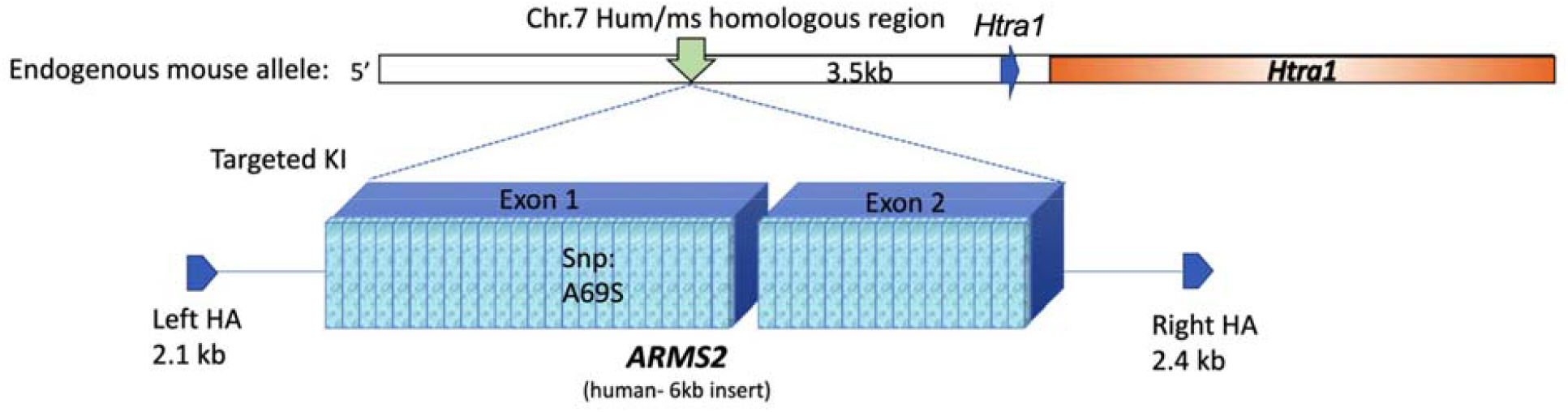
Targeting strategy for generation of the humanized *ARMS2*^*A69S*^ knock-in mouse. A 6kb human genomic fragment including the full gene sequence of *ARMS2* (exons 1 and 2; blue boxes); carrying the A69S substitution (rs10490924) in exon 1, and >2kb sequence upstream of *ARMS2* coding, was introduced at the region of homology between humans and mice on Chromosome 7, by homologous recombination, directed by 2.1-kb left and 2.4-kb right homology arms (HA). The *ARMS2*^*A69S*^ was positioned 3.5 kb upstream of *Htra1* (shown by green arrow). The transcriptional start site for *Htra1* is shown by a blue arrow.

### Tissue expression

To assess the expression and distribution of the inserted *ARMS2* gene in our expressed in our mouse model, we surveyed eight tissues from *ARMS2*^*A69S*^ mice and B6 controls by qRT-PCR (Figure 2). The *ARMS2* transcript was undetectable across all tissues in B6 mice, confirming the specificity of the assay for the human transgene. In *ARMS2*^*A69S*^ mice, the expression was markedly enriched in a subset of tissues, being highest in brain (mean ≈ 130-fold), followed by eye (≈ 50-fold) and testis (≈ 40 - 50-fold), with low-level to no expression in kidney, spleen, liver, adipose, and lung. These results were visualized by gel staining of RT-PCR products; a distinct *ARMS2* band was present in the *A69S* lanes for eye, brain, and testis (with fainter product in kidney, adipose and lung) and absent from spleen and liver, whereas all B6 lanes were negative; *Act-b* amplification was uniform across lanes, indicating comparable input. Together, these data show that transgene (Tg) expression, while detectable in several tissues, is concentrated in neural tissues including the eye.

**Figure 2:**
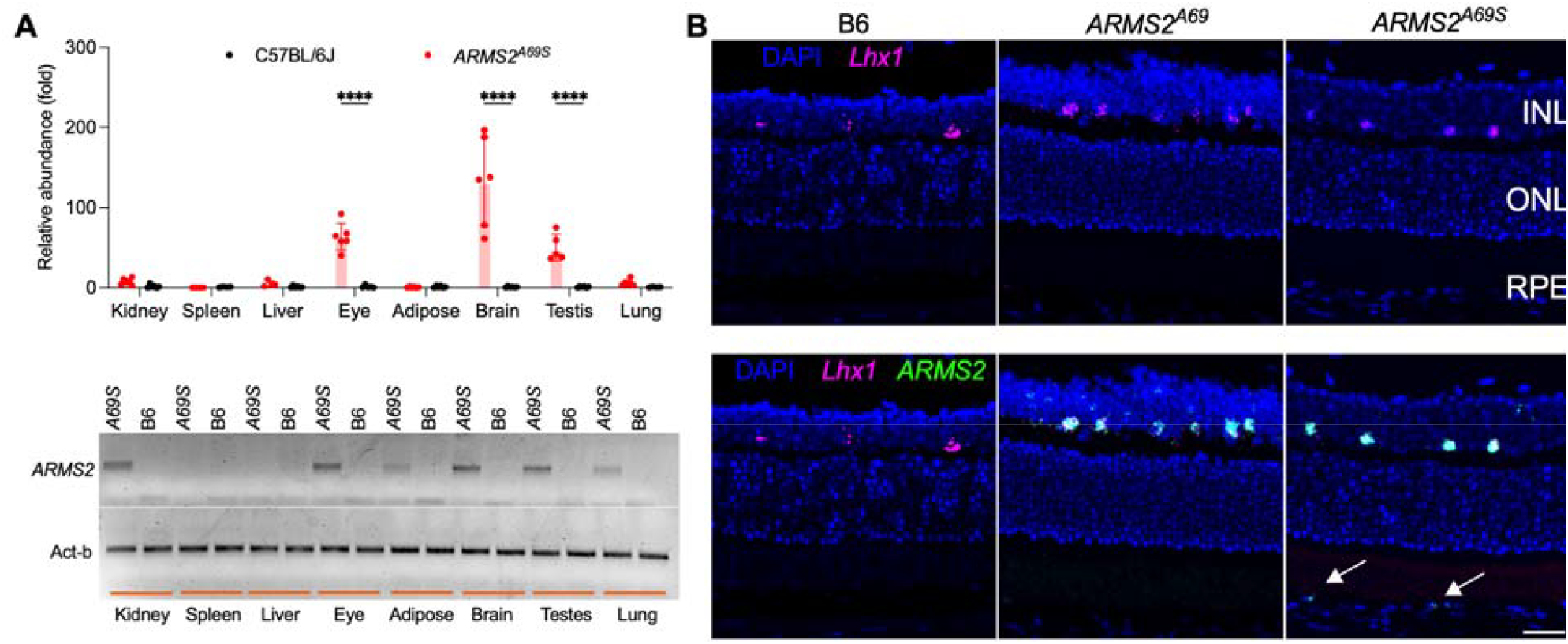
*ARMS2* expression. **A**. Relative *ARMS2* expression in *ARMS2*^*A69S*^ mice, as compared to B6. n = 4–6/tissue/strain (Top panel). A representative gel depicting *ARMS2* RT-PCR products in several *ARMS2*^*A69S*^ tissues is shown (Bottom panel). Tissue-specific *Actb* PCR products are shown as a loading control. **B**. RNA scope shows the expression of *ARMS2* (green) in horizontal cells (*Lhx1* marker; magenta) and RPE cells (white arrows) in the *ARMS2*^*A69S*^ and *ARMS2*^*WT*^ mice. n = 6/strain.

Commercially available ARMS2-specific antibodies tested either gave non-specific bands both in B6 and *ARMS2*^*A69S*^, or did not detect any Tg-specific bands in *ARMS2*^*A69S*^ tissues (not shown). Therefore, to localize *ARMS2* transcripts within the retina, we performed fluorescence in situ hybridization on cryosections from eyes of 6-month-old B6, *ARMS2*^*WT*^, and *ARMS2*^*A69S*^ mice (Figure 2B). In both *ARMS2*^*WT*^ and *ARMS2*^*A69S*^ retinas, *ARMS2* transcripts (green) were detected as discrete puncta in the INL, colocalizing with *Lhx1* transcripts, a marker for the horizontal cells in the inner nuclear layer (INL). *ARMS2*^*WT*^ and *ARMS2*^*A69S*^ transcripts were also detected in the RPE. No *ARMS2* signal was detected by RNAscope in B6 retina or RPE.

### Disease phenotypes observed by fundus phenotyping, OCT and ERG

To test if disease phenotypes were observable in aging *ARMS2*^*A69S*^ mice, fundus imaging was performed at 12 months of age. Uniformly sized, white fundus spots were seen in both *ARMS2*^*A69S*^ and *ARMS2*^*WT*^ mice with reduced frequency in B6 mice, however, spot number was variable in the strains (Figure 3A). Upon quantification, a higher number of spots was observed in *ARMS2*^*A69S*^ mice as compared to B6 controls. An increase in spots was observed in *ARMS2*^*A69S*^ compared to *ARMS2*^*WT*^ mice but was not significant. OCT was used to measure thicknesses of different retinal layers (Figure 3B). A small but significant (approx. 3.5%) decrease in outer nuclear layer (ONL) thickness was observed in the *ARMS2*^*A69S*^ mice as compared to *ARMS2*^*WT*^ and B6 controls in both dorsal and ventral regions of the eye. The *ARMS2*^*WT*^ mice did not show a decrease in the ONL, suggesting an A69S specific effect (Figure 3C). We also noted a significant increase in the inner nuclear layer (INL) thickness in the *ARMS2*^*A69S*^ mice in the ventral region. Consistent with the observed thinning of ONL, electroretinography showed a significant decrease in scotopic a- and b-wave amplitudes in the *ARMS2*^*A69S*^ mice, compared to the B6 mice (Figure 3D), indicating a decreased photoreceptor function in these mice.

**Figure 3:**
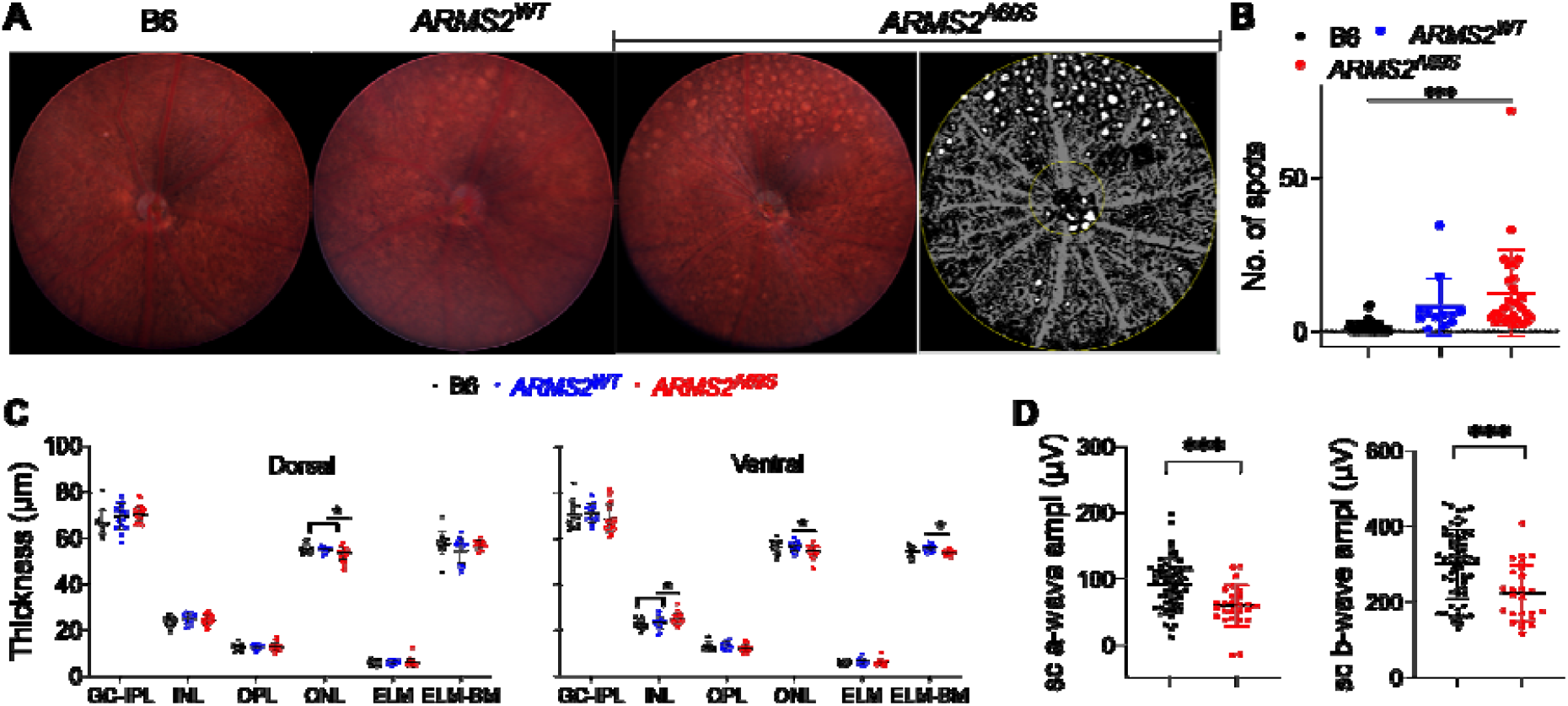
**A**. Fundus imaging shows the presence of spots in 12-month-old *ARMS2*^*A69S*^ mice. **B**. Quantification of spots using WeKa segmentation (right most image in panel A.) in Fiji using background, blood vessels or spots as three classifiers. n = 12-40/strain. **C**. Different retinal layers measured using optical coherence tomography (OCT), show a significant decrease in ONL thickness in 12-month-old *ARMS2*^*A69S*^ mice, compared to *ARMS2*^*A69S*^ and B6 mice, in dorsal and ventral regions. n = 14–22/strain. \**p* < 0.05. GC-IPL, ganglion Cell-inner plexiform layer, INL, inner nuclear layer, OPL, outer plexiform layer, ONL, uuter nuclear layer, ELM, external limiting membrane, BM, Bruch’s membrane. **D**. ERG shows a significant decrease in both scotopic a-(left) and b-wave amplitudes (right) in 12-month-old *ARMS2*^*A69S*^ mice. n = 26-58/strain. \*\*\**p* < 0.001.

### Transcriptomic analysis

To identify early expression changes before disease phenotypes are observable, an exploratory bulk RNA sequencing (RNA-seq) was performed on female retina and RPE-enriched tissue fractions obtained from postnatal day (P)14 mice. A total of 5,437 genes in the retina (4,011 down, 1,426 up) and 52 (15 down, 37 up) genes in the RPE were found to be differentially expressed in the *ARMS2*^*A69S*^ females, as compared to the B6 females (Figure 4A). Interestingly, we found that *Htra1* and *Plekha1*, the two genes that flank *ARMS2* on human Chromosome 10q26, to be upregulated in the RPE enriched fractions of *ARMS2*^*A69S*^, but not in the retina-enriched fractions. Of the 50 Hallmark pathways from the Molecular Signatures Database (MSigDB), gene set enrichment analysis identified 15 significantly enriched gene sets in retina and 9 in RPE (FDR < 0.05; Figure 4B). In retina, since the topmost significantly depleted set was mitochondrial (mt)–encoded oxidative phosphorylation (OXPHOS) genes (NES −1.99), we examined all the mt-encoded OXPHOS genes. At the level of individual mitochondrial genes (Figure 4C), reduction of mitochondrial genome–encoded OXPHOS transcripts in retina was broad, with 10 of 12 detected genes significantly decreased (log_2_FC =1.28 to −2.19) across Complexes I, III, and IV. Factors governing mitochondrial transcription were coordinately reduced in retina — *Tfam* (log_2_FC −0.84, padj 0.007), *Tfb2m* (−0.87, padj 0.011), *Tfb1m* (−0.68, padj 0.008) and *Mterf1a* (−0.95, padj 0.016). In RPE, the reduction was smaller and confined to a subset: *mt-Nd2, mt-Nd4* and *mt-Nd6* (Complex I) and *mt-Cytb* (Complex III) in which the reduction was approximately 30% (log_2_FC −0.47 to −0.56), whereas *mt-Co2, mt-Co3, mt-Atp6* and *mt-Atp8* showed non-significant increases (log_2_FC +0.69 to +1.59). In RPE, mitochondrial genome–encoded OXPHOS (NES −1.89) and mTORC1 signaling (−1.72) were depleted. Epithelial–mesenchymal transition was the most strongly enriched set (NES +2.41; z +8.95). Interestingly, we also found Peroxisome as one of the Hallmark gene sets to be significantly enriched in the retina. On building custom modules for the peroxisomal genes, the modules showed a downward trend in both retina and RPE tissues, suggesting the possibility of peroxisomes being affected in *ARMS2*^*A69S*^ mice (Figure 4D).

**Figure 4.**
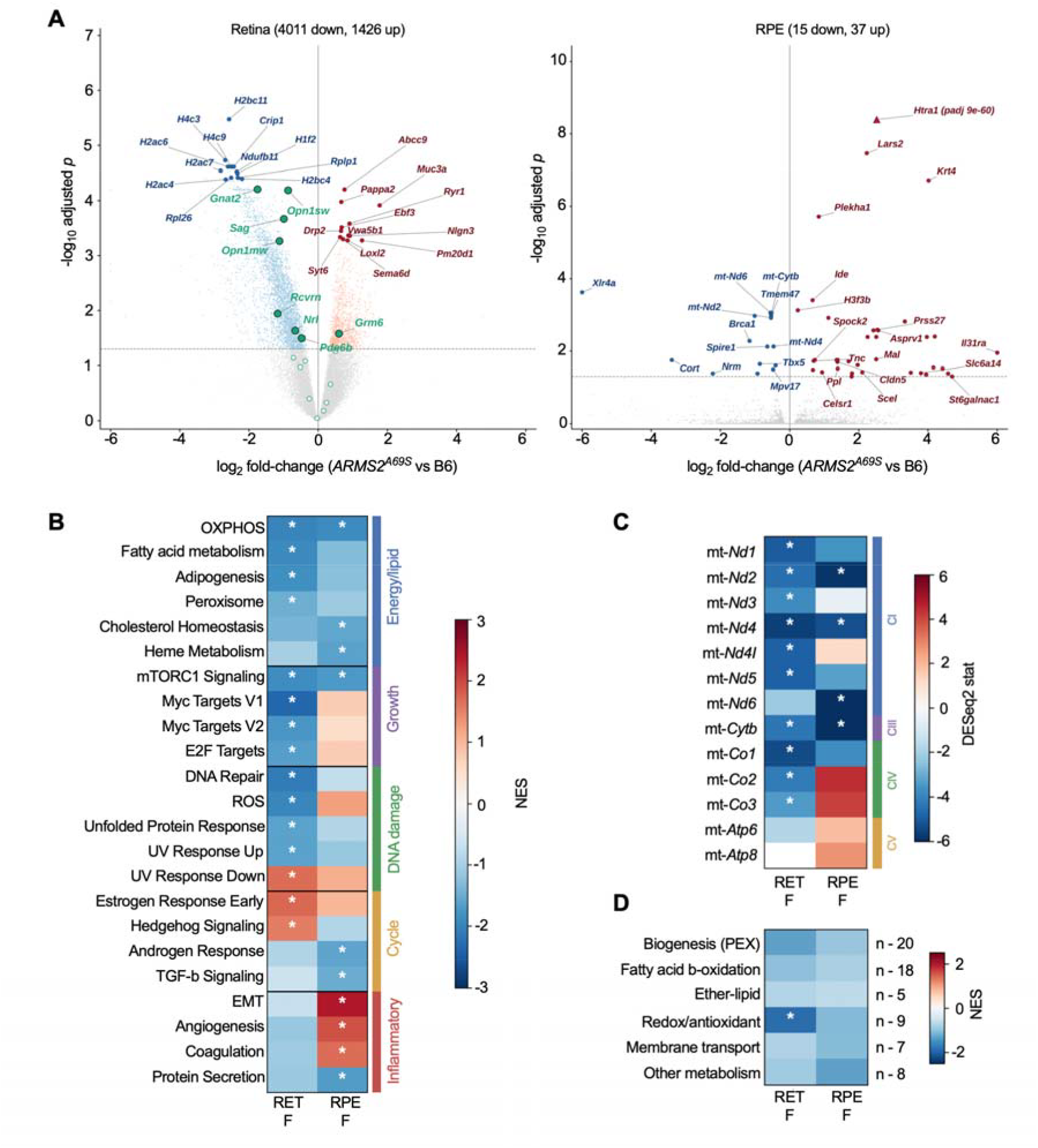
Transcriptomic effects of *ARMS2*^*A69S*^ in female retina and RPE. **A.** Volcano plots of differential expression between *ARMS2*^*A69S*^ and B6J retina (left) and RPE (right) showing up- and down-regulated genes. Log_2_-fold changes and adjusted *p*-values were obtained from DESeq2, the dashed line marks adjusted *p* = 0.05. **B**. Gene set enrichment analysis of Hallmark pathways significant in at least one tissue, grouped by biological theme, **C**. Per-gene effects for the 13 mitochondrial genome–encoded OXPHOS subunits, grouped by respiratory chain complex (CI, CIII, CIV, CV). Colour denotes the DESeq2 Wald statistic; asterisks denote adjusted *p* < 0.05, **D**. Gene set enrichment analysis of six curated peroxisomal functional modules; module sizes are shown at right. NES, normalized enrichment score from preranked GSEA (genes ranked by DESeq2 Wald statistic; 1,000 permutations), where positive and negative values indicate coordinated up- and down-regulation of a gene set in *ARMS2*^*A69S*^, compared to B6.

### mtDNA copy number change

To determine if the significant decrease in the transcriptome of the mitochondrial machinery observed in female retina was also reflected as a change in the mitochondrial content in *ARMS2*^*A69S*^ mice, we quantified relative mtDNA copy number in retina from *ARMS2*^*A69S*^, *ARMS2*^*WT*^, and B6 mice using NUMT-free mtDNA (mt-*Rnr2*) primers normalized to the nuclear *Tert* gene (Figure 5A). Relative mtDNA copy number was decreased significantly in the *ARMS2*^*A69S*^ mice compared to *ARMS2*^*WT*^, and also lower, but not significantly, when compared to B6 mice. A significant effect of sex was not seen, so both male and female samples within the individual strains were grouped together.

**Figure 5.**
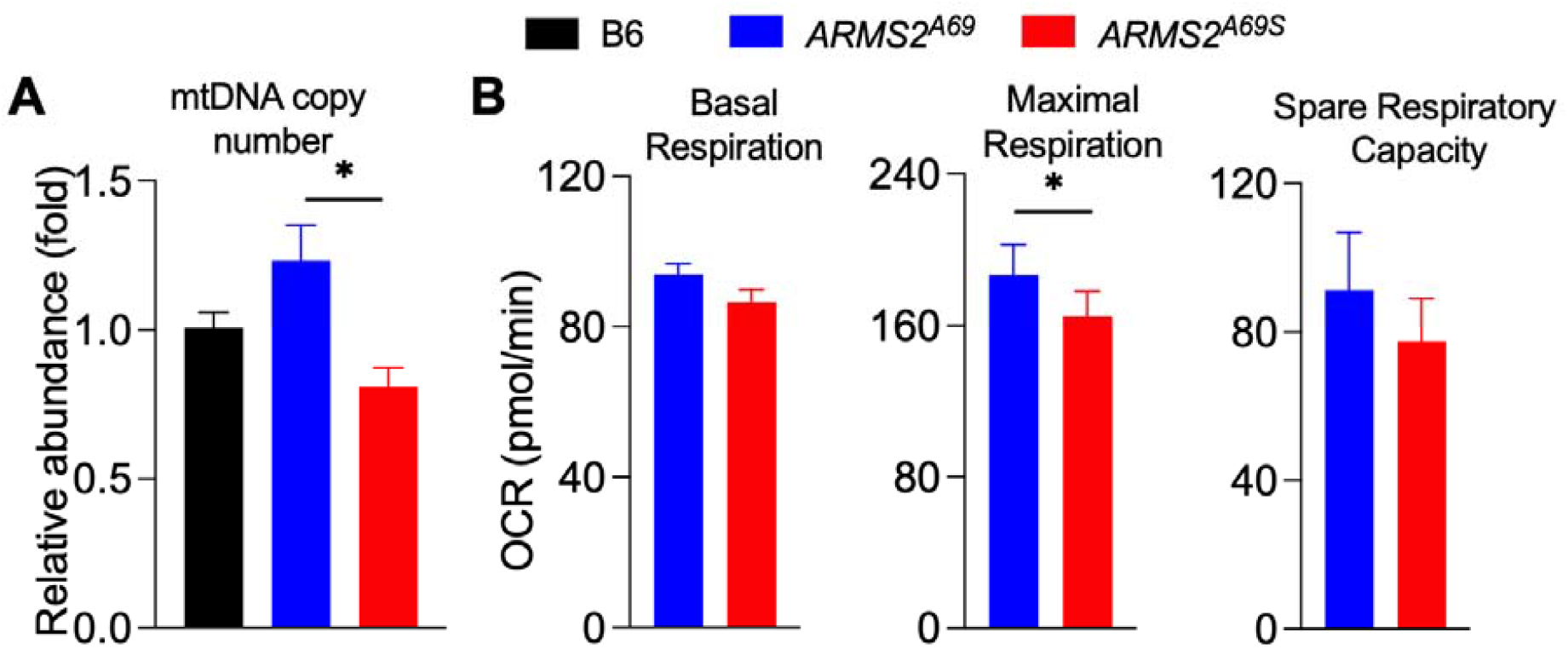
Mitochondrial copy number and functional changes noted in *ARMS2*^*A69S*^ mice. **A.** A decrease in mtDNA was noted in 3-month-old *ARMS2*^*A69S*^ using non-NUMT and n-*TERT* genes to calculate mtDNA:nDNA, n = 8 mice/strain, * *p* < 0.05. **B**. A decrease in OCR measurements noted in the retinal punches of 6-month-old old *ARMS2*^*A69S*^ as compared to the *ARMS2*^*A69*^ by Seahorse Mito Stress Test assay, n = 8 mice/strain, 3 punches/mouse.

### Mito Stress Test assay

To test the respiratory effect of the A69S variant, oxygen consumption rate (OCR) parameters were compared between *ARMS2*^*A69S*^ and its humanized wild-type control *ARMS2*^*WT*^ (Figure 5B). Using mixed-effects model analysis, maximal respiration was significantly lower in *ARMS2*^*A69S*^ than in *ARMS2*^*WT*^ (166.1 versus 187.7 pmol/min; difference = -21.7 pmol/min; F_1,5_ = 7.32, *p* = 0.043). The reduction was directionally consistent in both sexes (females, 175.7 versus 194.7 pmol/min; males, 156.4 versus 180.8 pmol/min), with no significant effect of sex (F_1,5_ = 0.30, p = 0.61) or strain-by-sex interaction (F_1,5_ = 0.11, *p* = 0.75). Spare respiratory capacity followed the same directional change but did not reach significance (78.5 versus 92.1 pmol/min; difference = -13.7 pmol/min; F_1,5_ = 1.35, *p* = 0.30). Basal respiration did not differ between strains (87.4 versus 90.9 pmol/min; F_1,5_ = 1.64, *p* = 0.26). Neither parameter showed a significant sex or interaction effect (all *p* > 0.29).

### MSI analysis

As lipid dysregulation in both the retina and RPE is characteristic of AMD, we examined the lipid content of 6-month-old RPE from *ARMS2*^*A69S*^, *ARMS2*^*WT*^, and B6 mice by mass spectrometry imaging (MSI). We first assessed the ordered trend for the main lipid classes in our preliminary MSI analysis, which suggested a statistically significant reduction in sphingomyelin (SM), such that the class-level SM intensity was lowest in *ARMS2*^*A69S*^ and lower in *ARMS2*^*WT*^, compared to B6 mice. Both sexes showed same directionality of change. However, we noticed a global intensity difference between some replicates. To ensure that the significant differences observed were not due to the global difference in signal intensity, we expressed each sphingolipid subclass as a share of the total sphingolipid pool to remove the global signal difference. This approach yielded statistically significant differences in the directional changes in lipid levels (Table 1). This correction reveals that the SM share falls while HexCer and Cer shares rise, and CerP remain flat. Our analysis suggests the pool is redistributed rather than simply reduced, which points toward altered flux through sphingomyelinase or glucosylceramide synthase (Table 1).

**Table 1.**
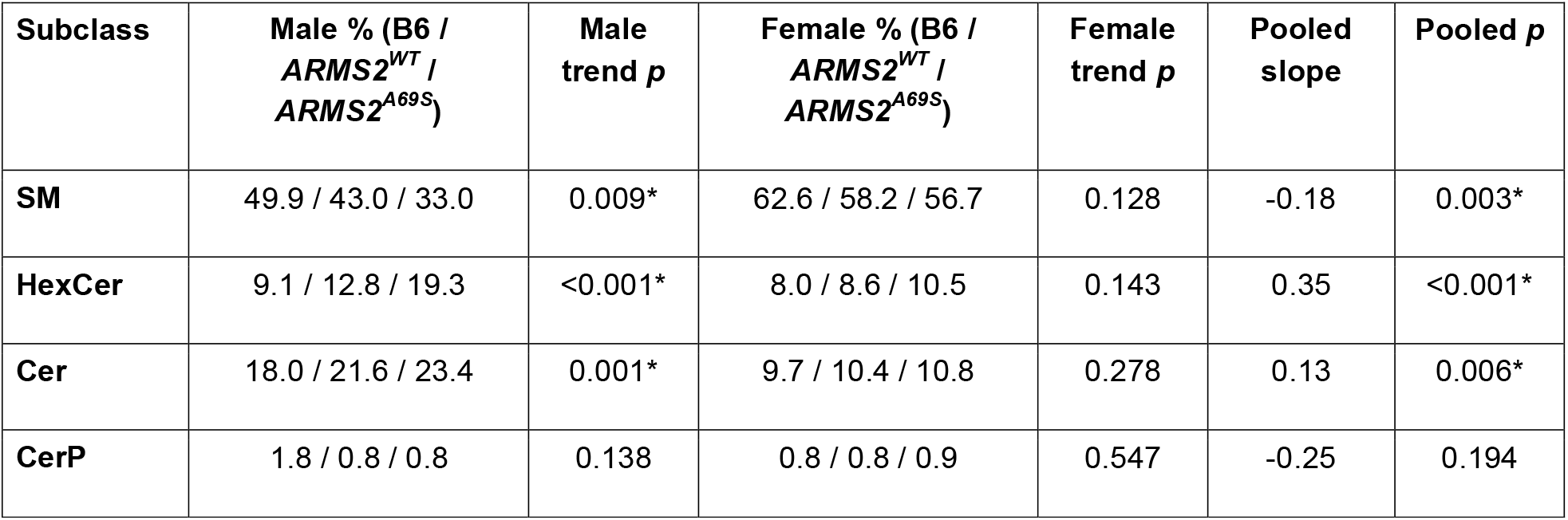
Sphingolipid subclass share of total sphingolipid signal (group means, %, percentage of individual subclass/total sphingolipid) and ordered-trend statistics. Shares are means of per-section percentages; trend p from linear contrast on log_2_share; * indicates q < 0.05.

## Discussion

We generated mouse models expressing a human *ARMS2*^*A69S*^ variant associated with increased risk of AMD and a control *ARMS2*^*WT*^strain, which carries the sequence found in non-affected individuals. Our characterization of these models suggests the constructs faithfully recapitulate the tissue-specific expression of *ARMS2* observed in humans, and reveals a subtle age-related phenotype that includes retinal degeneration and functional loss. Furthermore, changes in mRNA transcripts and lipid levels indicate a potential functional defect underlying mitochondrial dysfunction and altered lipid metabolism. The chief obstacle to establishing causality at the Chromosome 10q26 locus is that *ARMS2* and *HTRA1* are in strong linkage disequilibrium, so human association data cannot separate their contributions to AMD risk. Since *ARMS2* is not present in mice, we strategically placed our insertion of the full length *ARMS2*^*A69S*^ in a region of homology between humans and mice, immediately upstream of *Htra1*. This placement was intended to preserve the native cis-regulatory relationship the two genes to allow phenotypes to be attributable to the insertion in its native genomic context. Comparing these mice with *ARMS2*^*WT*^ KI mice allowed us to separate the effects due to the A69S variant from those of inserting the human *ARMS2* gene into the mouse genome.

### *ARMS2* is expressed in a tissue- and cell-type specific pattern which is not altered by the A69S mutation

Our qRT-PCR results confirmed the expression of the human *ARMS2* gene in the *ARMS2*^*A69S*^ mice, in a tissue-specific manner. Due to the lack of a suitable ARMS2 antibody, we used RNAscope *in situ* hybridization to identify the localization of *ARMS2* within the posterior eye. We confirmed that the A69S mutation did not alter the cell-type specificity in expression of *ARMS2* in the posterior eye as both *ARMS2*^*WT*^ and *ARMS2*^*A69S*^ mice showed *ARMS2* expression in horizontal cells of the inner retina as well as RPE cells. While no study has reproducibly validated the localization of *ARMS2* at the tissue and cellular level, our results are consistent with the human expression data presented in the Human Protein Atlas, where *ARMS2* is shown to be expressed in the testis, eyes and brain^43^. This confirms that the insertion of *ARMS2* in mice in the region of homology between mouse and humans may result in an expression pattern that is closer to humans.

### *In vivo* phenotypic changes noted in *ARMS2*^*A69S*^ mice

Since AMD is an aging disease, we characterized 12-month-old mice. Fundus imaging, OCT and ERG results confirmed that *ARMS2*^*A69S*^ mice develop disease phenotypes including fundus spots, decreased ONL thickness, and reduced scotopic a- and b-wave amplitudes, indicating degenerative effects on the PR cells. While *ARMS2* was not observed in the photoreceptor cells by RNAscope, it is plausible that the pathological changes in the RPE and horizontal cells lead to secondary effects in the PR cells. Immunohistochemistry with cell-specific markers will resolve if the fundus spots correspond to microglia or lipid accumulation or other abnormalities.

### Omics analysis suggests altered mitochondrial genome, which is supported by a change in mitochondrial number and function in the retina of *ARMS2*^*A69S*^ mice

Our initial transcriptomic analysis suggested a broad suppression of mitochondrial respiratory, peroxisomal, growth and stress-response programs in the female *ARMS2*^*A69*^ retina, along with the downward trend in RPE samples. The coordinate reduction of *Tfam, Tfb2m and Tfb1m* alongside mitochondrial and nuclear respiratory transcripts in female retina is consistent with reduced activity of the PGC-1α–NRF1–TFAM mitochondrial biogenesis program ^44-47^. Further, we saw a decrease in mitochondrial copy number and OCR measurements. Given that TFAM is a mtDNA packaging factor, and a direct determinant of copy number, a reduced expression of *Tfam* could result in a lower mitochondrial copy number; a lower level of mtDNA was observed in our analysis. We also observed a significant decrease in maximal and reduced respiratory spare capacity, without a change in basal OCR by the Seahorse Mito Stress test assay. Taken together, these results suggest a bioenergetic reserve deficit arising from the *ARMS2* AMD-risk allele, without a resting failure. The tissue meets baseline energy requirements, but a reserve deficit is a vulnerability state - normal under standard conditions, decompensating under stress, ageing, or oxidative load that is associated with late-onset, stress-dependent disease like AMD^48,49^.

### Lipidomic dysregulation in *ARMS2*^*A69S*^ RPE

Our MSI findings suggested a reduction of sphingomyelin species in enriched RPE of *ARMS2*^*A69S*^ mice and a shift in sphingolipid composition toward hexosylceramide and ceramide. Sphingolipids are central to RPE stress biology and have been implicated in AMD^50,51^. Ceramide generation drives RPE and photoreceptor apoptosis in oxidative-stress and light-damage models, and inhibition of ceramide synthesis is protective^52-55^. Sphingomyelin is the most abundant sphingolipid of the outer retina and RPE^51^, and its turnover by acid sphingomyelinase influences autophagic and phagolysosomal function in the RPE: lysosomal cholesterol accumulation secondary to lipofuscin bisretinoids activates acid sphingomyelinase and disrupts autophagy^56^, whereas loss of the enzyme results in AMD^57^. Hexosylceramide and ceramide increase in oxidatively stressed RPE cells^58^, and serum hexosylceramide is elevated in patients with late-stage AMD^59^. The simultaneous fall in SM share and rise in HexCer and Cer share is most consistent with increased catabolism of the sphingolipid pool rather than a change in its synthesis. Elevated sphingomyelinase activity would convert SM to ceramide (raising Cer share), with a portion of that ceramide channeled onward by glucosylceramide synthase to hexosylceramide (raising HexCer share); impaired lysosomal catabolism of glycosphingolipids could produce a similar redistribution. In contrast, a primary defect in SM synthesis would lower SM share without raising HexCer share, and so cannot by itself account for the pattern we observe. Both catabolic routes converge on the lysosomal–autophagic model of RPE dysfunction in AMD^60^. Moreover, given our evidence of altered mitochondrial biogenesis and function, together with the suggested mitochondrial localization of ARMS2^20,61^, both routes also implicate mitochondrial oxidative stress as an upstream driver of RPE injury in AMD^62^.

### Limitations and Future directions

We found an RPE-specific upregulation of *Htra1* expression in *ARMS2*^*A69S*^ mice in our transcriptomics analysis, with no change in expression levels in the retina. To resolve the effects of altered *Htra1* levels, a floxed *Htra1* mouse model might be used to study retina or RPE-specific knock-out effects on AMD-like phenotypes. Our transcriptomics analysismay be improved with a larger sample size for both retina and RPE-enriched samples that includes tissue from both male and female mice. Also, since mtDNA copy number alone cannot distinguish fewer mitochondria from fewer genomes per mitochondrion, a mass marker (TOMM20 or VDAC1 by western, or citrate synthase activity) might be used to assess whether reduced biogenesis or reduced nucleoid content accounts for mitochondrial dysregulation. Additional analysis using mitochondrial dyes like MitoView (Biotium) and TEM may confirm any decrease in mitochondrial numbers and alterations in mitochondrial structure. Absolute quantification by LC-MS and species identification by MS^2^ will be conducted to confirm the MSI results.

## Conclusion

Visual loss is a devastating consequence of AMD, significantly impacting the quality of life in the elderly. The *ARMS2*^*A69S*^ mouse model described here shows early mitochondrial changes. Since AMD is a complex disease, influenced by the interaction of multiple factors like genetics, age and environment, it is possible that this mitochondrial alteration due to the A69S genetic mutation in *ARMS2* increases the propensity of these mice to develop AMD-like phenotypes as they age and/or are faced with increased environmental stressors. Using this mouse model to understand how mitochondrial changes contribute to AMD-like features will be a crucial step in unravelling the disease mechanisms involved in AMD. This model will be valuable for elucidating the molecular and pathological mechanisms that results from the *ARMS2*^*A69S*^ variant, and to develop and test improved therapeutic strategies for degenerative eye diseases in the aging population.

## Acknowledgements

We gratefully acknowledge the contributions of Mass Spectrometry and Protein Chemistry Service (mass spectrometry imaging), Genome Technology (transcriptome sequencing) and Genomic Engineering Technology (generation of humanized mouse model) at JAX for expert assistance with the work described in this study.

## Data Availability

The data supporting the findings of this study are available within the article. Raw sequencing and mass spectrometry imaging datasets will be deposited in appropriate public repositories and released upon peer-reviewed publication. Additional data are available from the corresponding author upon reasonable request.

